# MiR-16-1^*^ and miR-16-2^*^ possess strong tumor suppressive and anti-metastatic properties in osteosarcoma

**DOI:** 10.1101/412411

**Authors:** Vadim V. Maximov, Saleh Khawaled, Zaidoun Salah, Lina Jaber, Nataly Bengaiev, Ahlam Barhoum, Marco Galasso, Eylon Yavin, Rami I. Aqeilan

## Abstract

Osteosarcoma (OS) is an aggressive malignancy affecting mostly children and adolescents. MicroRNAs (miRNAs) play important roles in OS development and progression. Here we found that miR-16-1* and miR-16-2* “passenger” strands as well as the “lead” miR-16 strand possess strong tumor suppressive functions in human OS. We report different although strongly overlapping functions for miR-16-1* and miR-16-2* in OS cells. Ectopic expression of these miRNAs affected primary tumor growth, metastasis seeding, and chemoresistance and invasiveness of human OS cells. Loss-of-function experiments verified tumor suppressive functions of these miRNAs at endogenous levels of expression. Using RNA immunoprecipitation (RIP) assays, we identify direct targets of miR-16-1* and miR-16-2* in OS cells. Furthermore, validation experiments identified *FGFR2* as a direct target for miR-16-1* and miR-16-2*. Overall, our findings underscore the importance of passenger strand miRNAs in osteosarcomagenesis.

**Novelty and Impact:** Osteosarcoma (OS) can be a fatal disease. MicroRNAs (miRNAs) play crucial roles in osteosarcomagenesis. In this study, we identify miR-16-1* and miR-16-2* as strong tumor suppressors and anti-metastatic genes in OS. This is the first report demonstrating tumor suppressive functions of passenger strands of these miRNAs in OS. Given that *MIR-16-1* is located in 13q14 region that is commonly deleted in several human malignancies, our findings shed light on oncogenic mechanisms triggered by 13q14 deletion.

## Introduction

Osteosarcoma (OS) is the most common form of bone and joint cancer which is responsible for about 9% of cancer deaths in children and adolescents ages 10-24 ^1^. Noteworthy, OS has a very complex karyotype with many nonrecurrent genetic abnormalities ^2^. Recently, it has been proposed that OS consists of many genetically different entities with different metastatic potential which occur due to OS-specific p53-independent preexisting genomic instability ^3^. Hence, genetic complexity of this malignant disease probably underlies the plateau in 5-year survival rate, which occurred in the middle of 1980s ^4^, with a third of all OS patients still dying during the first 5 years after diagnosis mainly due to pulmonary metastases ^5^. Moreover, metastases at diagnosis lead to further decrease in 5-year survival to less than one third of OS patients ^6-8^. Thus, better understanding of various osteosarcomagenesis molecular mechanisms is required for development of better treatments for this deadly malignant disease.

MicroRNAs (miRNAs) are small RNA molecules which length is typically 21-23 nucleotides although it may vary from 16 to 27 nucleotides ^9^. They mostly regulate protein-coding genes expression at the post-transcriptional level through mRNA translation repression and consequent degradation ^10^. Most of miRNA genes are transcribed by DNA-dependent RNA polymerase II and primary transcripts (pri-miRNAs) are further processed by the microprocessor complex in miRNA precursors (pre-miRNAs) with hairpin stem-loop structure, which are exported from the nucleus to the cytoplasm. Then, cytoplasmic RNase III Dicer cleaves pre-miRNAs which results in miRNA duplexes. Typically, one strand of a miRNA duplex is bound by argonaute proteins, loaded on miRNA-induced silencing complex (miRISC), and guides the miRISC to target mRNAs. This strand is called “lead” or “guide” strand. The other strand is usually mostly degraded and presented in the cell at a much lower level. This strand is called “passenger” or “star” strand and designated as miR* ^11^.

miRNAs expression deregulation as a driving force in oncogenesis is an established concept supported by massive research data ^12^. Involvement of miRNAs particularly in osteosarcomagenesis is also supported by accumulating evidences ^13^. Our group have identified a set of miRNAs, which expression is altered in OS samples in comparison with healthy bones ^14^. We further demonstrated that miR-27a/miR-27a* pair promotes OS metastasis at least partly through targeting of *CBFA2T3* ^15^. Pro-metastatic properties of “passenger” miR-27a* strand, which is expressed at much lower level than the “lead” miR-27a strand, were of particular interest for us. Indeed, there is mounting evidence that so-called “passenger” miRNA strands are involved in oncogenesis ^16-21^. This prompted us to study a possible involvement of other “passenger” miRNAs in osteosarcomagenesis. Noteworthy, miR-16, which originates from two loci, *MIR-16-1* (chromosome 13) and *MIR-16-2* (chromosome 3) (^22^ and Fig 1A), in the human genome, is down-regulated in OS samples versus healthy bones ^14^. Although *MIR-16-1* and *MIR-16-2* loci both encode the same miR-16 “lead” strand they encode different “passenger” strands – miR-16-1* and miR-16-2*, respectively. Interestingly, we found that miR-16-2* expression is also down-regulated in OS samples versus healthy bones^14^. We, therefore, decided to test miR-16-1* and miR-16-2* functions in osteosarcomagenesis. Here, we report that miR-16-1*, miR-16-2* as well as miR-16 possess tumor suppressive and anti-metastatic functions in human OS cells.

**Figure 1.**
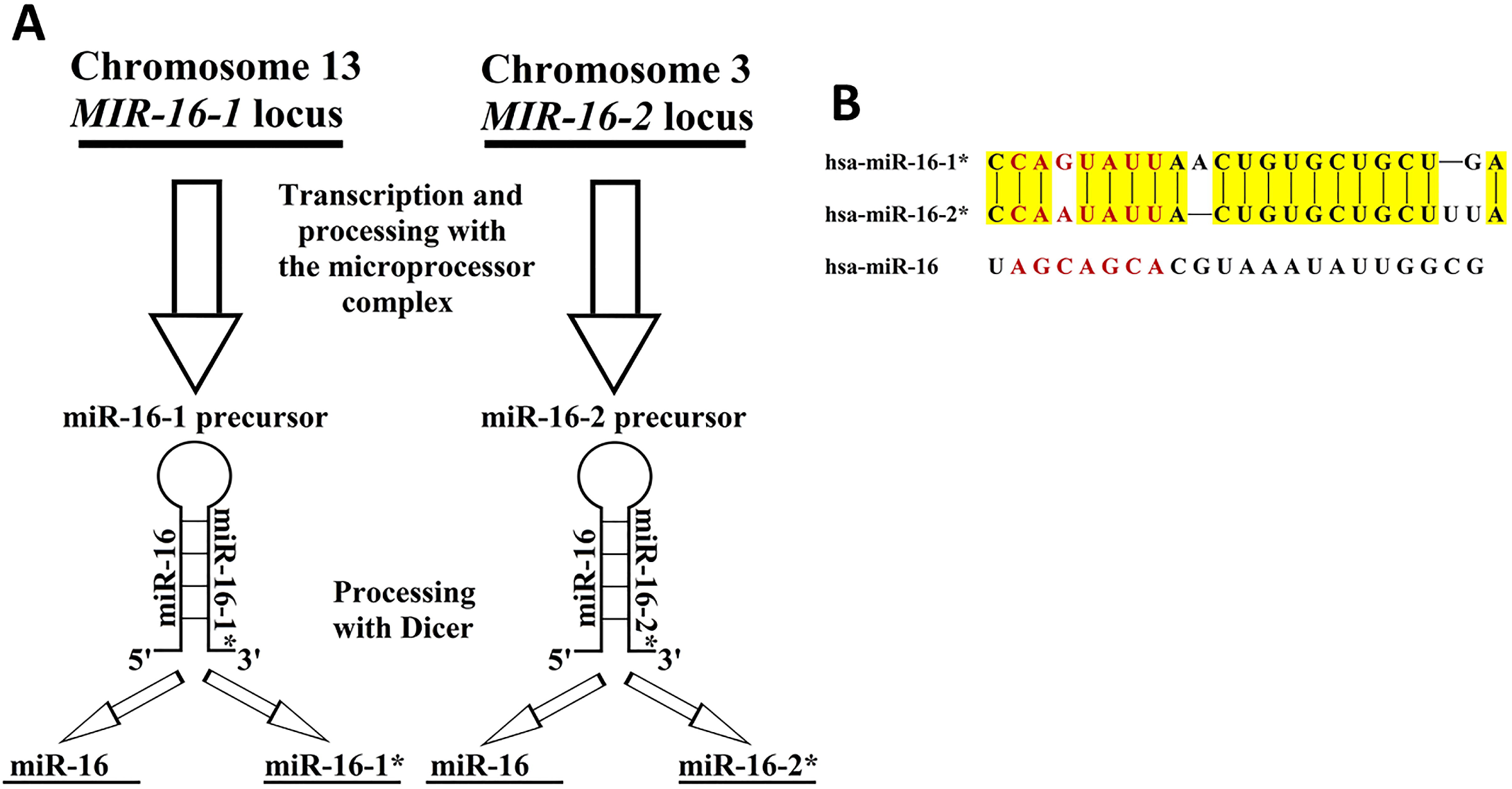
Scheme of genomic loci encoding uuman miR-16, miR-16-1* and miR-16-2*. (A) This is a schema depicting biogenesis of human miR-16, miR-16-1* and miR-16-2* from two different genomic *MIR-16* loci – *MIR-16-1* and *MIR-16-2*. (B) Similarity between sequences of human miR-16-1* and miR-16-2* is shown. Similarity regions are highlighted in yellow. Human miR-16 sequence is also shown for comparison. Red letters indicate seed regions of presented microRNAs. Error bars represent standard errors of mean.

## Materials and Methods

### Cell lines and cell culture

HOS, KHOS, U2OS, SAOS2, MG63, and 143B cells were obtained from American Type Culture Collection (ATCC, Manassas, VA) and were maintained in RPMI medium complemented with 10% heat-inactivated FBS, 100 U/ml penicillin, and 100 mcg/ml streptomycin. Where it is indicated in the text cells were cultured without antibiotics. HEK-293T cells were obtained from Fisher Scientific and were maintained in DMEM medium complemented with 10% heat-inactivated FBS, 2 mM L-glutamine, 100 U/ml penicillin, and 100 mcg/ml streptomycin.

### Colony formation assay

HOS or KHOS cells (1.5×10^2^) were seeded per a 10cm tissue culture plate in RPMI medium complemented with 10% heat inactivated FBS, 100 U/ml penicillin and 100 mcg/ml streptomycin. All experiments were conducted in four replicates. Colonies were fixed with ethanol, stained with Giemsa and counted in 9 days after seeding.

### Soft agar colony formation assay

6-well plates with 1.5 ml of 0.5% bottom agar in RPMI complemented with 10% heat-inactivated FBS, 2 g/L sodium bicarbonate, 2 mM L-glutamine, 100 U/ml penicillin, and 100 mcg/ml streptomycin were prepared by mixing even amounts of double concentrated RPMI containing double concentrated additives and 1% melted agar solution in water at 40°C. The bottom agar was left to solidify for 30 minutes at room temperature. Cells were added to prewarmed to 30°C double concentrated RPMI with double concentrated additives and mixed with even amount of cooled to 39°C 0.7% melted agar solution in water. The obtained 0.35% melted top agar solution in RPMI with additives containing cells was plated in prepared wells of 6-well plates with bottom agar. 1.5 ml of the top agar containing one thousand cells were plated in each well. 0.5 ml of RPMI medium with additives were added to each well after the top agar was solidified. Each experiment was conducted in triplicates. 0.2-0.3 ml of fresh RPMI medium with additives were added to each well twice a week in order to avoid drying of agar. Twenty one days after plating, the grown colonies in agar were fixed and stained with 0.01% crystal violet solution in 10% ethanol in water. Then wells were washed 6 times with water and colonies were counted.

### Chemoresistance assays

U2OS (10×10^3^) or HOS (5×10^3^) cells were plated in each well of 96-well plates. Cisplatin or doxorubicin treatments were started 24 hours after plating by complete replacing of medium in wells with 200 mcl of medium containing required concentration of the drug. Each drug concentration was tested in triplicates. Cells survival was quantified by an XTT-based assay (Biological Industries, USA) according to the manufacture’s protocol in 48 hours after the drug adding. Survival of untreated cells was taken as 100% for each type of cells.

For experiments with synthetic microRNA mimics, U2OS cells in a well of 6-well plates were transfected with 200 pmoles of a double-stranded synthetic microRNA mimic and 17.5 mcl of Lipofectamine 2000 transfection reagent (Thermo Fisher Scientific, USA) according to the manufacturer’s instructions. Cells were in RPMI medium with 10% heat-inactivated serum but without antibiotics and at 90% confluence. 6-8 hours after transfection the medium was changed for fresh one without antibiotics. Twenty four hours after transfection cells were seeded in wells of 96well plates (ten thousand cells per a well) and treatments with drugs were started 16 hours later.

### Matrigel invasion assay

HOS cells at 50-70% confluency were detached from culture plates by incubation in 1 mM EDTA in PBS. Two hundred thousand cells were placed in the upper part of a blind well chemotaxis chamber in serum-free RPMI medium. The bottom part of the blind well chemotaxis was filled with RPMI medium complemented with 10% heat-inactivated FBS, 100 U/ml penicillin, and 100 mcg/ml streptomycin. Upper and bottom parts of the blind well chemotaxis chamber were separated by a Matrigel-coated membrane with size of pores 8 mcm. Invasion assays were conducted for 4 hours in triplicates. The upper surfaces of membranes were wiped with a cotton swab in order to remove non-invaded cells. Then invaded cells were fixed and stained with Diff-Quick System (Dade Behring, Inc., UK). Photos of ten random fields for each membrane were taken and invaded cells were counted.

### Tumorigenic and metastatic assays in NOD/SCID mice

All experiments with NOD/SCID mice were conducted in agreement with guidelines of the Institutional Animal Care and Use Committee of The Hebrew University of Jerusalem under approved protocols. Four-six weeks aged NOD/SCID male mice were used for experiments. One million HOS cells or five hundred thousand KHOS cells were subcutaneously (SC) injected in 100 mcl of RPMI medium in each flank of a NOD/SCID mouse. Each mouse was injected in both, right and left, flanks. Measurements of linear tumors’ sizes were conducted once or twice a week after tumors appeared. Tumors’ volumes were estimated as (a*b^2)/2 where a is the longest linear size and b is the smallest linear size. The experiment with HOS clones overexpressing miR-16, miR-16-1* or miR-16-2* was conducted for 23 days. The experiment with KHOS cells overexpressing miR-16 or miR-16-2* was conducted for 64 days. The experiment with KHOS cells overexpressing Contr-Sp, miR-16-Sp, miR-16-1*-Sp or miR-16-2*-Sp was conducted for 48 days. Tumors’ masses were measured when mice were euthanized and open. Pictures of fluorescent EGFP-positive metastatic nodules in lungs were taken in the experiment with EGFP-positive KHOS cells overexpressing microRNA sponges. Then numbers of metastatic nodules in each lung were counted.

Orthotopic intratibial (IT) injections were conducted in right rare legs of four-six weeks old NOD/SCID mice. Five hundred thousand HOS cells in 20 mcl of 25% growth factors-reduced matrigel in RPMI were injected each time. IT injected NOD/SCID mice were observed for 31 days. Then mice were euthanized and all measurements were conducted. Tumors’ volumes were estimated as (a*b^2)/2 where a is the longest linear size and b is the smallest linear size.

### Statistical analysis

Statistical significance of all pair-wise comparisons was assessed by two-or one-sided Student’s t-test. If data variability was too large for Student’s t-test then the Rank-Sum statistics was applied. Adjusted p-value was applied to estimate statistical significance of finding in the RIP-Seq experiment. P-value adjustment was conducted by the Benjamini-Hochberg method. Further details can be found in the Supporting Information for this article online. Statistical significance of the correlations investigated was estimated by Spearman Correlation.

**Other materials and methods can be found in the Supporting Information for this article online.**

## Results

### MiR-16, miR-16-1* and miR-16-2* expression in OS samples and OS cell lines

We found earlier that expression of miR-16 and miR-16-2* is downregulated in OS versus healthy bone while miR-16-1* expression was undetectable ^14^. All these miRNAs originate from two loci – *MIR-16-1* and *MIR-16-2* (Fig 1A). The lead miR-16 strand is identical for both loci while the miR-16-1* strand is specific for the *MIR-16-1* locus and the miR-16-2* strand is specific for the *MIR-16-2* locus (Fig 1A). MiR-16-1* and miR-16-2*, which have been mostly considered so far as passenger strands or miR-16 biogenesis byproducts, have very similar but somewhat different sequences (Fig 1B). There is also one mismatch between miR-16-1* and miR-16-2* sequences in the seed region (Fig 1B), which should lead to different binding sites for these miRNAs.

Nothing has been studied about functions of the passenger miR-16-1* and miR-16-2* strands, and we decided to address their possible roles in osteosarcomagenesis. First, we addressed abundance of these miRNAs in OS cells and assayed miR-16, miR-16-1* and miR-16-2* expression in several OS cell lines by TaqMan Real-Time PCR (Suppl Fig 1). MiR-16-1* expression was out of the quantitative range for this assay (Suppl Fig 1K). MiR-16 expression varied from 5.5 thousand to 28.5 thousand molecules per cell (Suppl Fig 1G), which is similar to earlier published data for mouse organs and embryo^23^. MiR-16-2* expression varied from ∼320 to ∼850 molecules per cell (Suppl Fig 1H) and the ratio of miR-16 expression to miR-16-2* expression was in the range from 18 to 80 (Suppl Fig 1J).

### MiR-16, miR-16-1* and miR-16-2* overexpression suppresses OS cells survival

In order to address functional roles of miR-16, miR-16-1* and miR-16-2* in OS we overexpressed these miRNAs in HOS cells. Since HOS polyclonal cultures lose overexpression of these miRNAs too quick (data are not shown) we obtained HOS clones for overexpression of each miRNA (Fig 2A).

**Figure 2.**
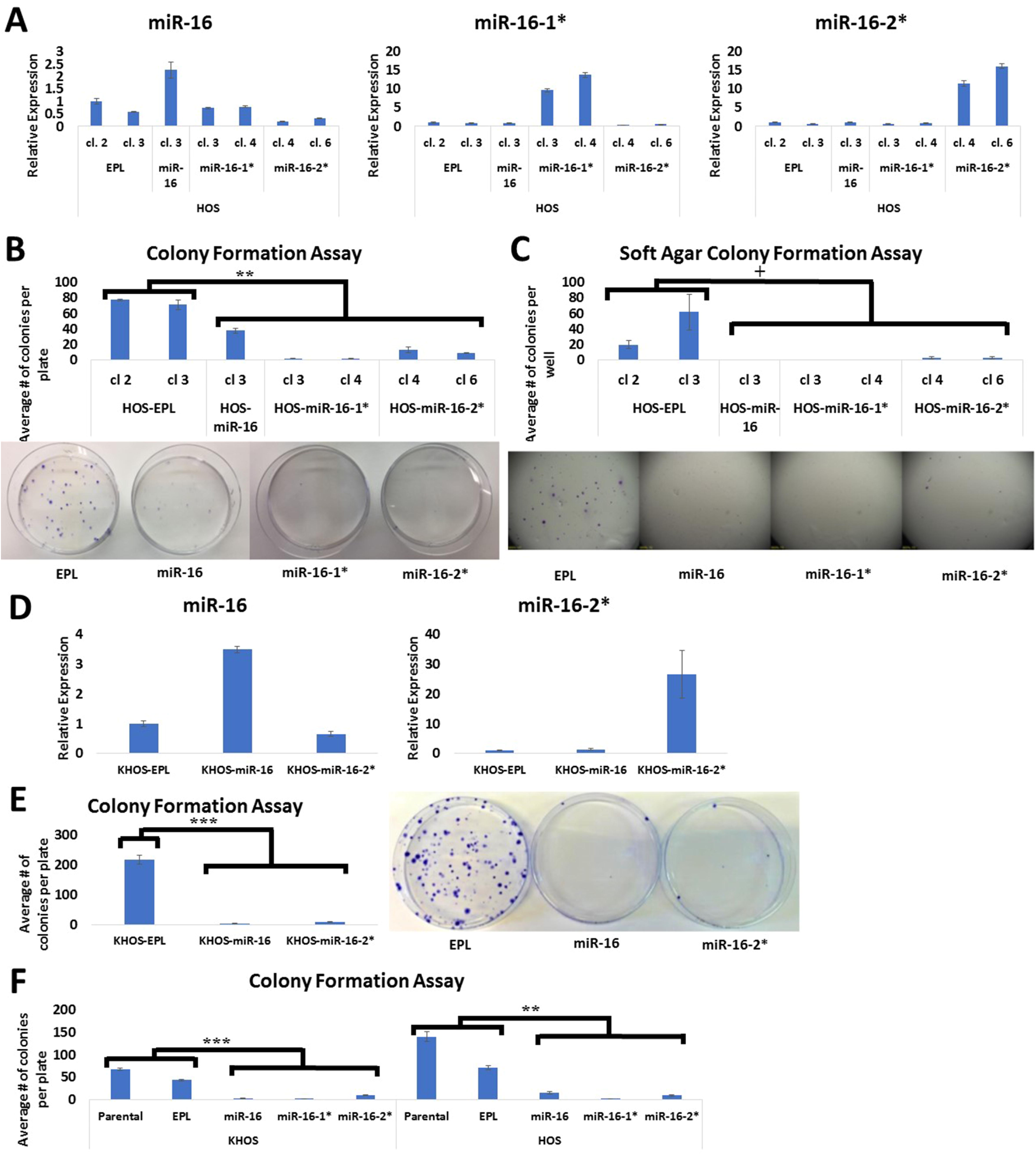
Effects of human miR-16, miR-16-1* and miR-16-2* overexpression on OS cells survival and anchor-independent growth. (A) Overexpression of miR-16, miR-16-1* and miR-16-2* in HOS clones was assessed by SYBR green-based Real-Time PCR. Expression of microRNAs in each sample was normalized to U44 expression. MicroRNAs’ expression in the sample HOS-EPL, cl. 2 was taken as one. (B) A colony formation assay for HOS clones was conducted in four replicates in 10cm plates for each clone. One hundred fifty cells were seeded per each 10cm plate. Top – quantification of the assay; Bottom – representative pictures. (C) A soft agar colony formation assay for HOS clones was conducted in triplicates in wells of 6-well plates for each clone. One thousand cells were plated per a well of 6-well plate. Top – quantification of the assay; Bottom – representative pictures. (D) Overexpression of miR-16 and miR-16-2* in KHOS cells was assessed by SYBR green-based Real-Time PCR. Expression of microRNAs in each sample was normalized to U44 expression. MicroRNAs’ expression in the sample KHOS-EPL was taken as one. (E) A colony formation assay for KHOS cells was conducted in four replicates in 10cm plates for each type of cells. One hundred fifty cells were seeded per each 10cm plate. Right – quantification of the assay; Left – representative pictures. (F) KHOS and HOS cells were infected with lentiviruses for overexpression of corresponding microRNAs at MOI=200. The same KHOS cells were applied in experiments shown on Figures 3A and 3B (F) Each type of KHOS and HOS cells was subjected to the colony formation assay three days after infection. The colony formation assay was conducted in four replicates in 10cm plates for each type of cells. One hundred fifty cells were seeded per each 10cm plate. EPL stays for empty puromycin lentivirus. Student’s t-test was applied everywhere with the exception of the figure (C) in order to estimate statistical significance. Rank-sum statistics was applied in the figure (C) in order to estimate statistical significance. Error bars represent standard errors of mean. ** – two-sided p-value □ 0.01 for Student’s t-test; *** – two-sided p-value □ 0.001 for Student’s t-test; + – two-sided p-value ≤ 0.05 for Rank-sum statistics.

Results of colony formation and soft agar colony formation assays suggest that overexpression of any of these miRNAs – miR-16, miR-16-1*, or miR-16-2* – drastically reduces both colony formation ability and anchor-independent growth of HOS cells (Fig 2B, C). Noteworthy, effects of miR-16-1* as well as miR-16-2* overexpression on HOS cells colony formation is much stronger than the effect of miR-16 overexpression (Fig 2B). These data suggest that miR-16, miR-16-1* as well as miR-16-2* overexpression decreases HOS cells survival.

Further we decided to check whether miR-16, miR-16-1* and miR-16-2* overexpression effects can be reproduced in another OS cell line. We chose KHOS cell line and were able to overexpress there miR-16 and miR-16-2* (Fig 2D) but not miR-16-1* (data are not shown). Overexpression of miR-16 and miR-16-2* led to a dramatic decrease in KHOS colony formation ability (Fig 2E), which is consistent with our data for HOS cells (Fig 2B). We hypothesized that our inability to overexpress miR-16-1* was due to a very strong effect of miR-16-1* on KHOS cells survival and selection with puromycin kills KHOS cells overexpressing miR-16-1*. In order to overcome this obstacle, we subjected KHOS cells, which were infected with lentiviruses for overexpression of miR-16, miR-16-1* or miR-16-2*, to colony formation assays immediately after infection without puromycin selection. Indeed, KHOS cells infected with miR-16, miR-16-1* or miR-16-2* had much reduced colony formation abilities in comparison to KHOS cells infected with empty lentivirus or parental KHOS cells (Fig 2F). Similar results were obtained for HOS cells (Fig 2F). Altogether, these findings suggest that products of *MIR-16-1* and *MIR-16-2* loci have anti-survival effects in vitro. These findings are consistent with possible tumor suppressive functions of these miRNAs.

### MiR-16, miR-16-1* and miR-16-2* overexpression enhances apoptosis and chemosensitivity

We also set to determine whether these miRNAs promote apoptosis of OS cells. Indeed, overexpression of miR-16, miR-16-1* or miR-16-2* increases the percentage of annexin V/PI staining suggesting that KHOS cells are undergoing apoptosis (Fig 3A, B).

**Figure 3.**
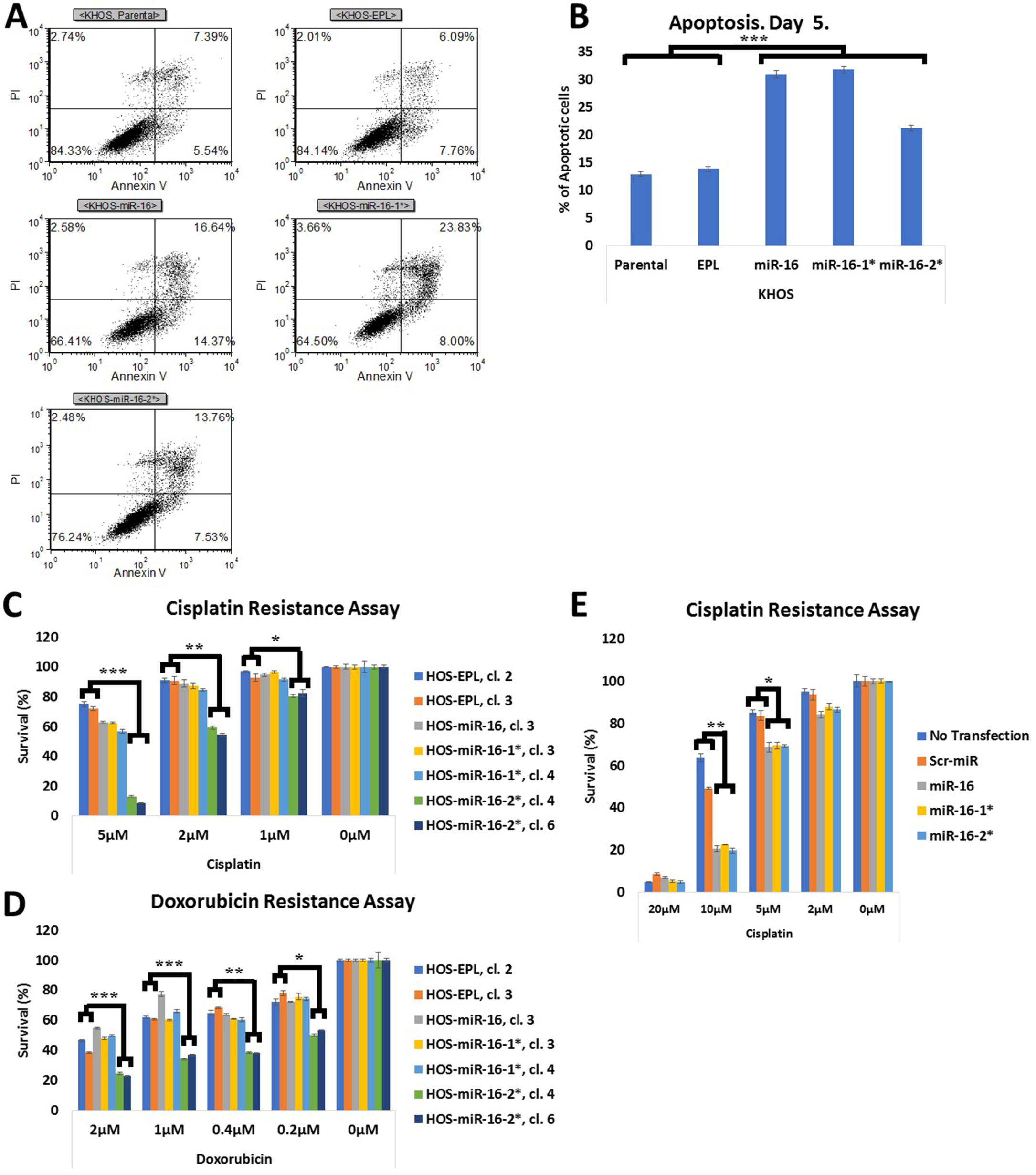
Effects of human miR-16, miR-16-1* and miR-16-2* overexpression on OS cells apoptosis and chemoresistance. (A) and (B) KHOS cells were infected with lentiviruses for overexpression of corresponding microRNAs at MOI=200. The same KHOS cells were applied in the experiment shown on Figure 2F. Annexin V-FITC/PI staining was conducted for KHOS cells five days after infection. (A) Representative flow cytometry dot plots are shown. (B) Quantification of the apoptotic assay is shown. Cells in the right bottom quadrant were considered as early apoptotic cells and cells in the right top quadrant were considered as late apoptotic cells. Total percentages of apoptotic cells are shown on the histogram. (C, D) HOS clones were subjected to cisplatin and doxorubicin resistance assays as described in “Materials and Methods”. Cytotoxicity was quantified by the XTT assay. (E) U2OS cells were transfected with 200 pmols of a corresponding synthetic microRNA mimic. Ten thousand cells were seeded for the cisplatin resistance assay in 16 hours after transfections in wells of a 96-well plate. Cisplatin was added in 24 hours after seeding and the cisplatin resistance assay was conducted as described in “Materials and Methods”. Cytotoxicity was quantified by the XTT assay. EPL stays for empty puromycin lentivirus. Student’s t-test was applied in order to estimate statistical significance. Error bars represent standard errors of mean. * – two-sided p-value L 0.05 for Student’s t-test; ** – two-sided p-value □ 0.01 for Student’s t-test; *** – two-sided p-value □ 0.001 for Student’s t-test.

We next addressed the outcome of these miRNAs on chemosensitivity. To this end, we assessed effects of miR-16, miR-16-1* and miR-16-2* overexpression on HOS resistance to cisplatin and doxorubicin treatment. Remarkably, miR-16-2* overexpression but not miR-16-1* or miR-16 overexpression significantly sensitized HOS cells to both cisplatin and doxorubicin treatments (Fig 3C, D). Curiously, the same miR-16-1* overexpressing clones in parallel colony formation assays formed even less number of colonies than corresponding miR-16-2* overexpressing clones (Fig 2B). These data suggest that despite of high similarity between miR-16-1* and miR-16-2* sequences these miRNAs possess different although overlapping functions.

We also addressed the question whether synthetic miR-16, miR-16-1* and miR-16-2* mimics could affect OS cells resistance to cisplatin. We chose U2OS cell line for these experiments since this cell line is easy to transfect. Interestingly, all tested microRNAs – miR-16, miR-16-1* and miR-16-2*, sensitized U2OS cells to the cisplatin treatment with similar efficiencies (Fig 3E). This result suggests that at high level of overexpression of all tested miRNAs – miR-16, miR-16-1* and miR-16-2*, can efficiently sensitize OS cells to the cisplatin treatment. It also suggests that miR-16, miR-16-1* and miR-16-2* mimics could be beneficial to improve outcomes of the conventional OS chemotherapy, which includes combined treatment with cisplatin, doxorubicin and methotrexate ^5^.

### Endogenous miR-16, miR-16-1* and miR-16-2* expression effects OS cells chemoresistance *in vitro*

In order to clarify whether miR-16, miR-16-1* and miR-16-2* endogenous expression effects chemosensitivity of OS cells we constructed miR-16, miR-16-1* and miR-16-2* sponges [SP] (Fig 4A) and overexpressed them in U2OS cells. Upon overexpression, miRNA sponges bind the corresponding miRNAs and sequester them from binding with their targets. Thus, miRNA sponges’ overexpression leads to functional inactivation of corresponding miRNAs ^24^. We subjected U2OS cells, which overexpress miR-16, miR-* or miR-16-2* sponges, to the cisplatin and doxorubicin resistance assays. While none of these sponges had a consistent effect on doxorubicin resistance (data are not shown) overexpression of any of these sponges increased the resistance of U2OS cells to cisplatin (Fig 4B).

**Figure 4.**
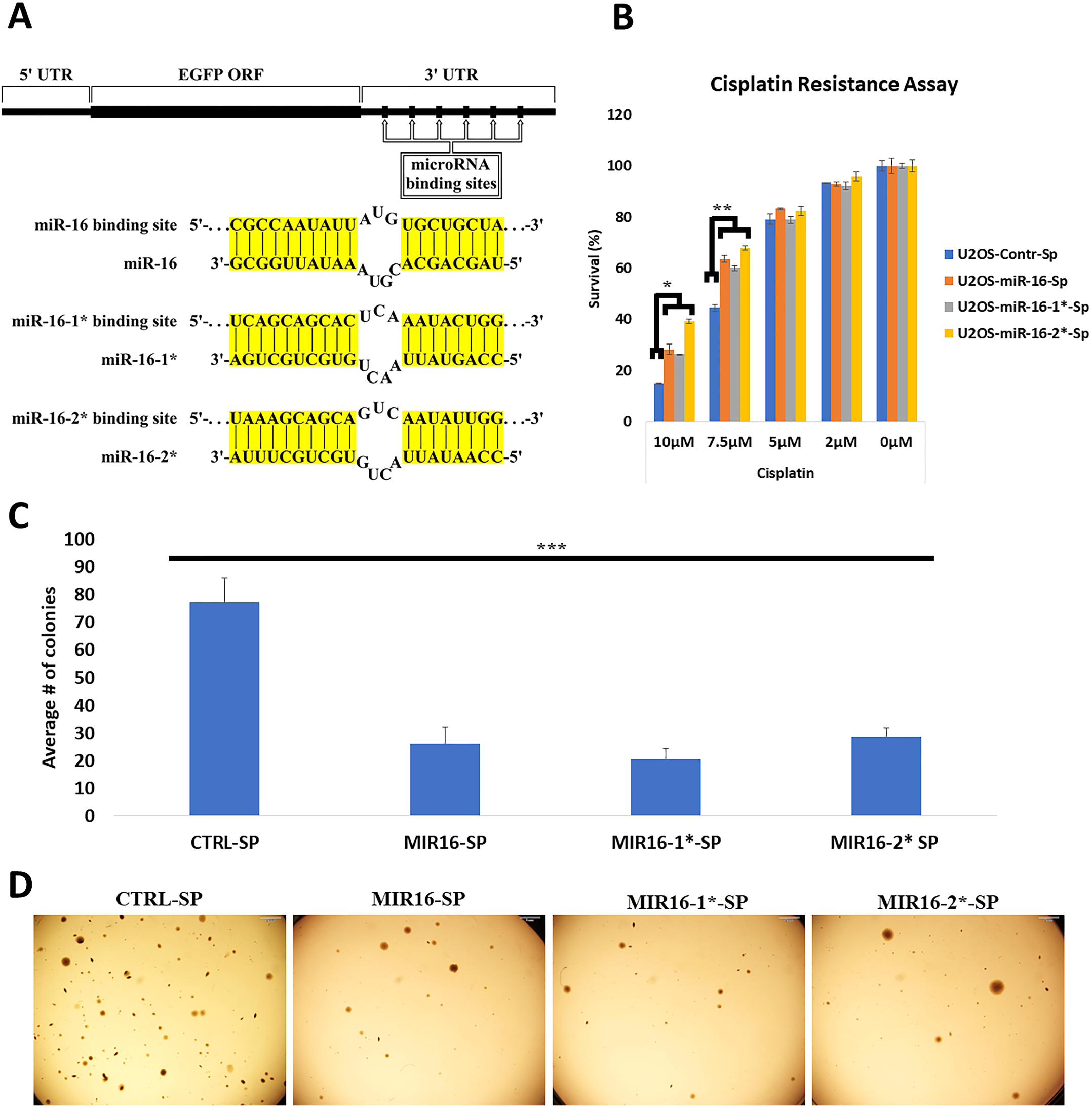
Effects of human miR-16, miR-16-1* and miR-16-2* sponges on U2OS cells cisplatin resistance. (A) A microRNA sponge is schematically depicted on the top of this picture. MiR-16, miR-16-1* and miR-16-2* binding sites and complementary interactions of these binding sites with corresponding microRNAs are shown below. Regions of complementary interactions between microRNAs’ binding sites and corresponding microRNAs are highlighted in yellow. (B) U2OS cells overexpressing microRNAs’ sponges were subjected to the cisplatin resistance assay as described in “Materials and Methods”. Cytotoxicity was quantified by the XTT assay. Student’s t-test was applied in order to estimate statistical significance. Error bars represent standard errors of mean. * – two-sided p-value □ 0.05 for Student’s t-test; ** – two-sided p-value □ 0.01 for Student’s t-test.

We also conducted CRISPR-based knock-out of each *MIR-16* locus separately and both of them together in U2OS cells (Suppl Fig 2). Knock-out of any of two *MIR-16* loci separately increased resistance of U2OS cells to cisplatin and knock-out of both *MIR-16* loci increased resistance of U2OS cells to cisplatin treatment even stronger (Suppl Fig 2E). These results are consistent with microRNA sponges data (Fig 4B), and suggest that endogenous miR-16, miR-16-1* as well as miR-16-2* expression is essential for sensitivity of OS cells to cisplatin treatment.

### MiR-16, miR-16-1* and miR-16-2* overexpression effects on OS cells tumorigenesis in NOD/SCID mice *in vivo*

In order to assess effects of miR-16, miR-16-1* and miR-16-2* overexpression on the ability of human OS cells to form tumors in vivo, we subcutaneously injected cells from HOS clones overexpressing corresponding miRNAs into NOD/SCID mice. MiR-16-1* as well as miR-16-2* overexpression completely abolished the ability of HOS cells to form subcutaneous tumors in NOD/SCID mice while miR-16 overexpression significantly reduced HOS cells tumorigenesis upon their subcutaneous injection in NOD/SCID mice (Fig 5A-C). These results suggest that miR-16-1* as well as miR-16-2* possess stronger tumor suppressive activities than the lead miR-16 strand.

**Figure 5.**
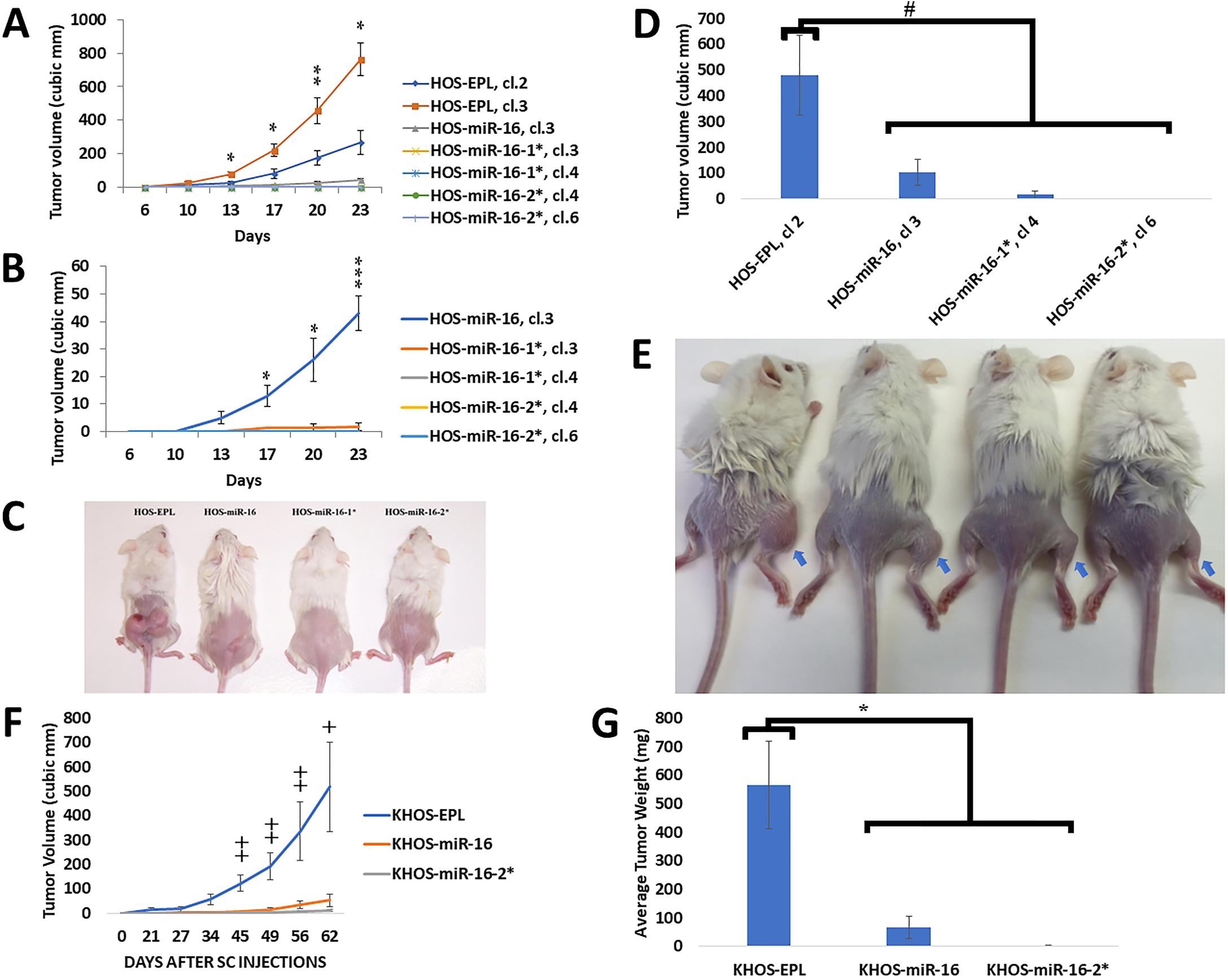
Effects of human miR-16, miR-16-1* and miR-16-2* overexpression on human OS cells tumorigenesis in NOD/SCID mice *in vivo*. (A)-(C) HOS clones overexpressing corresponding microRNAs were subcutaneously injected in NOD/SCID mice. Each HOS clone was injected in 5 NOD/SCID mice. Each mouse was injected in both, right and left, flanks (n = 10 per each HOS clone). HOS-EPL, cl. 2 and HOS-EPL, cl. 3, which are infected with the empty lentivirus, were applied as controls. (A) Time course of tumor growth for all HOS clones is presented. (B) Time course of tumor growth for HOS clones excluding the control HOS clones is presented. (C) Representative pictures of NOD/SCID mice with subcutaneous tumors are shown. (D),(E) NOD/SCID mice were injected in the right rare leg intratibially with HOS clones as described in “Material and Methods”. Five hundred thousand cells were used per each injection. Six mice were injected for each – HOS-EPL, cl. 2 (n=6) and HOS-miR-16-1*, cl. 4 (n=6) clones. Five mice were injected for each – HOS-miR-16, cl. 3 (n=5) and HOS-miR-16-2*, cl. 6 (n=5) clones. Mice were open and all measurements were conducted on day 31 after injections. (D) Final measurements of tumors’ volumes are shown. (E) Representative pictures of NOD/SCID mice with intratibial tumors are shown. Blue arrows indicate the injected leg. (F),(G) NOD/SCID mice were subcutaneously injected with KHOS cells overexpressing corresponding microRNAs. Each NOD/SCID mouse was injected in both, right and left, flanks. Five hundred thousand KHOS cells were used per each injection. KHOS-EPL cells, which are infected with the empty lentivirus, were applied as a control. Two mice were injected with KHOS-EPL cells (n=4). Four mice were injected with KHOS-miR-16 cells (n=8). Three mice were injected with KHOS-miR-16-2* cells (n=6). (F) Time course of tumor growth is presented. (G) Final measurements of tumors’ volumes are shown. Blue arrows indicate injected legs. Tumors’ volumes were measured as described in “Materials and Methods”. Subcutaneous injections were conducted as described in “Material and Methods”. EPL stays for empty puromycin lentivirus. Student’s t-test was applied everywhere with the exception of the figure (F) in order to estimate statistical significance. Rank-sum statistics was applied in the figure (F) in order to estimate statistical significance. Error bars represent standard errors of mean. * – two-sided p-value □ 0.05 for Student’s t-test; ** – two-sided p-value □ 0.01 for Student’s t-test; *** – two-sided p-value □ 0.001 for Student’s t-test; # – one-sided p-value □ 0.05 for Student’s t-test; + – two-sided p-value ≤ 0.05 for Rank-sum statistics; ++ – two-sided p-value □ 0.01 for Rank-sum statistics.

We also assessed the ability of selected HOS clones to form tumors in the orthotopic environment. MiR-16, miR-16-1* as well as miR-16-2* overexpression significantly reduced HOS tumorigenesis upon intratibial injections in NOD/SCID mice (Fig 5D, E). These results further support tumor suppressive properties of miR-16, miR-16-1* and miR-16-2* in OS cells *in vivo*.

We also verified tumor suppressive functions of miR-16 and miR-16-2* in another OS cell line – KHOS. KHOS cells overexpressing miR-16 as well as miR-16-2* formed significantly smaller tumors upon subcutaneous injections of these cells in NOD/SCID mice than the control KHOS cells infected with the empty lentivirus (Fig 5F, G). Unfortunately, we could not conduct the same experiment with KHOS cells overexpressing miR-16-1* since these cells could not survive selection with puromycin (see above). Nevertheless, these data further support that miR-16 as well as miR-16-2* possess tumor suppressive properties in OS cells.

### MiR-16, miR-16-1* and miR-16-2* overexpression effects on OS cells invasion *in vitro*

We also tested whether miR-16, miR-16-1* or miR-16-2* overexpression affects metastatic properties of OS cells such as invasion. Indeed, miR-16, miR-16-1* as well as miR-16-2* overexpression significantly reduced HOS cells invasion in vitro (Fig 6). This data suggests possible involvement these microRNAs in OS metastatic process.

**Figure 6.**
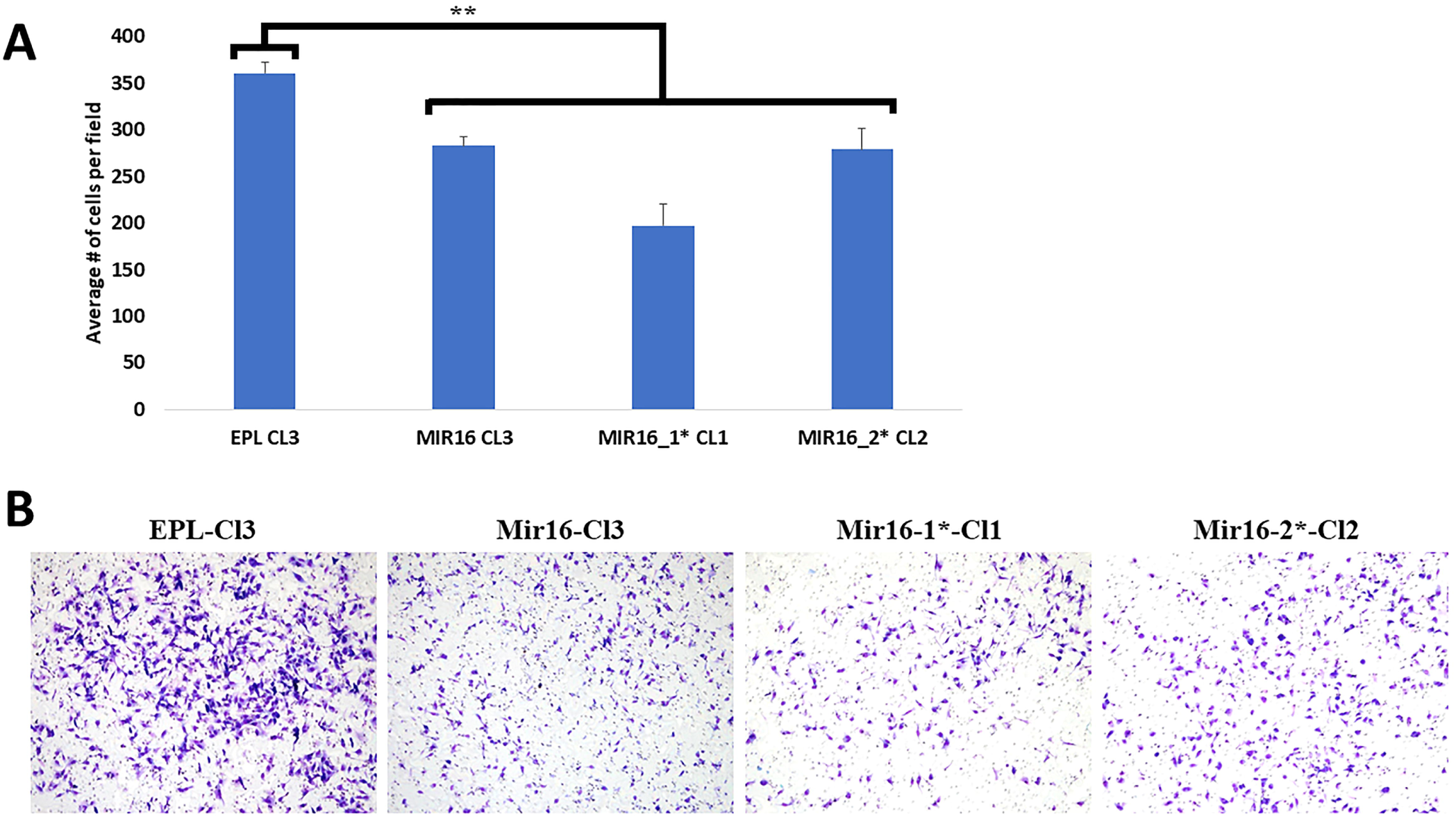
Effects of human miR-16, miR-16-1* and miR-16-2* overexpression on invasion of HOS cells *in vitro*. Trans-well invasion *in vitro* assay for HOS clones overexpressing corresponding microRNAs was conducted as described in “Material and Methods”. Two hundred thousand cells were applied per a well. This assay was conducted in triplicates for each HOS clone. The assay was stopped and the invaded cells were counted in 4 hours. HOS-EPL, cl. 3, which is infected with the empty lentivirus, was applied as a control. (A) Quantification of the invasion assay is presented. (B) Representative pictures of fields with invaded cells are shown. Student’s t-test was applied in order to estimate statistical significance. ** – two-sided p-value □ 0.01 for Student’s t-test.

### Roles of endogenous miR-16, miR-16-1* and miR-16-2* expression in OS cells tumorigenesis and metastatic process upon subcutaneous injections of human OS cells in NOD/SCID mice *in vivo*

Although overexpression data clearly indicated tumor suppressive functions of miR-16, miR-16-1* and miR-16-2* in OS cells it was unclear whether endogenous levels of these miRNAs affects tumorigenic and/or metastatic properties of OS cells. In order to clarify this question, we subcutaneously injected KHOS cells overexpressing sponges for these miRNAs into NOD/SCID mice. Overexpression of a sponge for any of these miRNAs led to an increase in the average primary tumor mass although it was not statistically significant (Suppl Fig 3A-C). Nevertheless, there was a significant increase in the number of lung metastases in KHOS cells overexpressing a sponge for any of these miRNAs in comparison to KHOS cells overexpressing the control miRNA sponge (Suppl Fig 3D,E). These data suggest that miR-16, miR-16-1* as well as miR-16-2* likely possess anti-metastatic activities in human OS cells *in vivo* at endogenous levels of expression.

### MiR-16, miR-16-1* and miR-16-2* direct targets in OS cells

The preceding observation indicate anti-survival, pro-apoptotic, tumor suppressive and anti-metastatic functions of miR-16, miR-16-1* and miR-16-2* in human OS cells. However, molecular mechanisms underlying these functions of miR-16, miR-16-1* and miR-16-2* remained obscure. We therefore aimed to identify direct targets of these miRNAs which is essential for clarifying of the molecular mechanisms.

In order to achieve this aim, we chose an unbiased approach based on immunoprecipitation of argonaute proteins’ complexes, known as RIP, for RNA immunoprecipitation (Suppl Fig 4). Argonaute proteins are an indispensable part of microRNA-induced silencing complexes (mRISCs), which contain direct targets of all miRNAs. Hence, RNA from immunoprecipitated argonaute proteins’ complexes should contain all direct targets of all miRNAs. We assumed that overexpression of a particular miRNA should recruit more of its direct targets into mRISCs and hence immunoprecipitation of the argonaute proteins’ complexes should identify those direct mRNA targets. The outline of this approach is shown in supplementary Figure 4.

In order to implement this approach, we transfected cells from HOS clones overexpressing corresponding miRNAs as well as control HOS cells with a mix of plasmids overexpressing FLAG-and MYC-tagged Ago1, Ago2, Ago3 and Ago4 proteins. Then argonaute proteins’ complexes were immunoprecipitated by the FLAG-tag and total RNA was purified from these immunoprecipitants. The purified total RNA was subjected to deep RNA sequencing. This experiment was conducted in biological triplicates for each microRNA (miR-16, miR-16-1* and miR-16-2*) and for the control HOS cells. Enrichment of every annotated RNA in the immunoprecipitated argonaute proteins’ complexes upon overexpression of each studied microRNA was evaluated (Suppl Table 1).

In order to highlight the most promising candidate direct targets of the studied miRNAs we reanalyzed miRNA and mRNA expression data from our previous article ^14^. Indeed, expression of direct targets of a given miRNA, which expression is mostly regulated by this miRNA in OS cells, should inversely correlate with expression of this miRNA in OS samples. Thus, we determined genes, which expression inversely correlates with miR-16 (Suppl Table 2) and/or miR-16-2* (Suppl Table 3) expression in OS samples at RNA level. Overlaps of these genes with genes, which are enriched in argonaute proteins’ immunoprecipitants for miR-16 (Suppl Table 4) and for miR-16-2* (Suppl Table 5), represent the most promising candidate direct targets for these miRNAs. We further refined our sets of candidate direct targets based on presence of predicted binding sites for corresponding miRNAs (Suppl Tables 4 & 5). Unfortunately, we could not conduct similar analysis for miR-16-1* targets, since its expression was not technically detectable by miRNA microarrays in most OS and healthy bone samples ^14^.

We chose 6 potential direct targets for miR-16 and 9 potential direct targets for miR-16-2* based on our analysis (Suppl Table 6). We also chose one specific potential direct target for miR-16-1* (Suppl Table 6). We checked expression of these targets in HOS clones overexpressing corresponding miRNAs. Most of these genes were significantly upregulated in HOS clones overexpressing corresponding microRNAs at levels comparable to their enrichment in the immunoprecipitated argonaute proteins’ complexes (data are not shown). These data suggested that these genes were enriched in the immunoprecipitated argonaute proteins’ complexes solely due to upregulation of their expression. However, expression of 3 genes (*FRAS1*, *MMP16* and *IFI6*) was not changed or changed weakly in HOS clones overexpressing corresponding miRNAs (Suppl Fig 5A-F). Thus, these genes are likely to be directly regulated at the level of translation by corresponding microRNAs.

Verification of potential direct targets of studied miRNAs by Real-Time PCR suggested that increase in proportion of argonaute bound mRNA upon a miRNA overexpression indicates that this mRNA is a direct target of the miRNA. Simple enrichment of a mRNA in argonaute immunoprecipitants upon a miRNA overexpression is frequently due to expression upregulation of this mRNA (data are not shown). Therefore, direct targets could be even among genes, which are depleted in argonaute immunoprecipitants upon a microRNA overexpression. In order to check this possibility, we made a list of targets, which expression is inversely correlated with miR-16-2* expression in OS samples and which are depleted in the immunoprecipitated argonaute proteins’ complexes upon miR-16-2* overexpression (Suppl Table 7). We chose *FGFR2* gene for a detailed study. *FGFR2* is significantly depleted in argonaute immunoprecipitants upon miR-16-2* overexpression (Suppl Tables 1, 7), *FGFR2* expression inversely correlates with miR-16-2* expression in OS samples (Suppl Tables 5, 7) and *FGFR2* has predicted binding sites for miR-16, miR-16-1* and miR-16-2* (Suppl Table 6). In addition, *FGFR2* is frequently activated and/or overexpressed in gastric cancer ^25^, a potential target for cancer therapy ^26^, involved in OS metastasis ^27^ and its mutations lead to inherited craniofacial malformation syndromes, which are associated with bone abnormalities ^25^. Interestingly, a proportion of argonaute-bound *FGFR2* mRNA is significantly increased in miR-16 overexpressing HOS cells and not changed in miR-16-2* overexpressing HOS cells (Suppl Fig 5G, H). However, Ct values for *FGFR2* from miR-16-2* overexpressing HOS cells were very high and the results were not quantitative. Hence, we cannot conclude whether the proportion of argonaute-bound *FGFR2* mRNA is increased in miR-16-2* overexpressing HOS cells. To validate whether *FGFR2* is a direct target of miR-16 and its associated passenger strands, we cloned its 3’UTR containing miRNA-responsive elements into the pGL3 vector downstream of the *firefly luciferase* open reading frame and assessed the reporter activity in control, miR-16-1*-overexpressing and miR-16-2*-overexpressing HOS clones. The luciferase activity of 3’UTR was markedly reduced by miR-16-1/2* overexpression as compared to control cells (Suppl Fig 5I). Moreover, miR-16, miR-16-1* as well as miR-16-2* overexpression leads to increase in Akt Ser473 phosphorylation (Suppl Fig 5J) that is consistent with PI3K/Akt pathway up-regulation in *FGFR2* overexpressing osteoblasts ^28^. Altogether, these results suggest that miR-16 passenger strands could have overlapping targets hence contributing to OS development and progression.

## Discussion

Here, we have provided evidences that miR-16-1* as well as miR-16-2* “passenger” strands function as tumor suppressors in OS. Their tumor suppressive effects as strong or even stronger than tumor suppressive effects of the “lead” miR-16 strand in human OS cells. These miRNAs effect both, OS primary tumorigenesis and metastasis. They have anti-survival and pro-apoptotic action in human OS cells and also reduce invasiveness and chemoresistance of human OS cells. This is the first report of any function for miR-16-1* and/or miR-16-2*. Although anti-metastatic properties of miR-16-1* in gastric cancer cells were reported earlier by Wang T and his co-workers ^29^, it was not sufficiently detailed and explored as in our study. In fact, Wang T *et al* overexpressed miR-16-1 precursor for all functional experiments and did not check for miR-16 overexpression ^29^. Since miR-16-1* precursor overexpression is expected to lead to overexpression of both, the ”lead” miR-16 and the “passenger” miR-16-1*, strands (Fig 2A), then, it is not clear which strand caused functional effects in gastric cancer cells ^29^. Noteworthy, our data also indicate that miR-16-1* and miR-16-2* have different although strongly overlapping functions. In addition, our results suggest that *FGFR2* is a direct target of miR-16-1* as well as miR-16-2*. This sheds some light on mechanisms underlying miR-16-1* and miR-16-2* tumor suppressive functions since *FGFR2* is frequently activated in gastric cancer ^25^, promotes metastasis in OS ^27^, its aberrant activation leads to inherited craniofacial malformation syndromes, which are associated with bone abnormalities, ^25^ and it regulates survival and differentiation of osteoblasts ^30^.

Dr. Croce’s group was the first to present evidence for tumor suppressive functions of miR-16 in attempts to explain a role of 13q14 deletions in chronic lymphocytic leukemia (CLL). They found *MIR-16-1* locus deletion and/or miR-16 down-regulation in about two-third of all CLL cases ^31^. Indeed, knock-out of *miR-16-1* locus in mice caused CD5-positive B-cell malignancies with penetrance of ∼25% ^32^. Curiously, knock-out of *miR-16-2* locus in mice caused CD5-positive B-cell malignancies with 100% penetrance ^33^. However, no reports of *MIR-16-2* locus deletion in human CLL can be found. One explanation may be a possible difference in the pattern of *MIR-16-1* and *MIR-16-2* loci expression between humans and mice. Nevertheless, all the mice data point toward tumor suppressive functions of both *miR-16-1* and *miR-16-2* loci. Noteworthy, no association between 13q14 deletion and miR-16 expression in human CLL was found^34^. Our data, although obtained in human OS cells, gives a strong indication that down-regulation the “passenger” miR-16-1* strand expression rather than the “lead” miR-16 strand may be behind oncogenic effects of the *MIR-16-1* locus deletion in CLL and other malignancies.

Although our data suggest strong tumor suppressive properties of miR-16, miR-16-1*, and miR-16-2*, obligatory knock-outs of neither *miR-16-1* nor *miR-16-2* loci were reported to cause osteosarcoma in mice ^32, 33^. It suggests that either additional oncogenic events are needed to cause OS in mice or miR-16, miR-16-1*, and miR-16-2* expression down-regulation is involved in later stages of osteosarcomagenesis in mice or both possibilities together. Differences in mechanisms of osteosarcomagenesis between mice and humans are also possible. Predominant involvement of miR-16, miR-16-1*, and miR-16-2* in later stages of osteosarcomagenesis, particularly, in OS metastasis and chemoresistance, seems to be highly possible in agreement with our data. Indeed, miR-16, miR-16-1*, and miR-16-2* affect mostly metastatic and chemoresistance properties of human OS cells at endogenous levels of expression.

Our data about effects of synthetic miR-16, miR-16-1*, and miR-16-2* mimics on chemoresistance of human OS cells *in vitro* suggest a potential use of these mimics to improve the outcomes of the conventional OS chemotherapy. Noteworthy, miR-34a and miR-16 mimics-based drugs are already in clinical trial for treatment of different malignancies (reviewed in ^3^).

## Acknowledgments

Authors thank Dr. Yuval Nevo (Bioinformatic division, Hebrew University-Hadassah Medical School, Jerusalem, Israel) for bioinformatic analysis of RIP experiments.

This research work was funded by a grant from the Liddy Shriver Sarcoma Initiative (R.I.A.) and Yissum seed funds (R.I.A. & E.Y). V.V.M. was supported by ICRF post-doctoral grant, Lady Davis fellowship and Israel Ministry of Immigrant Absorption postdoctoral fellowships.

## Abbreviations

OS: osteosarcoma
miRISC: microRNA-induced silencing complex
mRNA: messenger RNA
CLL: chronic lymphocytic leukemia
RIP: RNA Immunoprecipitation

